# Gsx2 but not Gsx1 is necessary for early forebrain patterning and long-term survival in zebrafish

**DOI:** 10.1101/2021.09.13.460150

**Authors:** RA Coltogirone, EI Sherfinski, ZA Dobler, SN Peterson, AR Andlinger, LC Fadel, RL Patrick, SA Bergeron

**Affiliations:** Department of Biology, West Virginia University, Morgantown, WV, USA; Department of Neuroscience, West Virginia University, Morgantown, WV, USA

**Keywords:** zebrafish, neurodevelopment, transcription factor, forebrain patterning, gs homeobox

## Abstract

Central nervous system (CNS) development is regulated by regionally expressed transcription factors that impart initial cell identity, connectivity, and function to neural circuits through complex molecular genetic cascades. *genomic screen homeobox 1* and *2* (*gsx1* and *gsx2*) encode homeobox transcription factors expressed in the developing CNS in multiple vertebrates examined to date. However, we have limited knowledge of the expression of these transcription factors and the gene networks that they regulate across developing brain regions in zebrafish. The objective of this study was to comprehensively examine *gsx1* and *gsx2* expression throughout neurodevelopment and characterize *gsx1* and *gsx2* mutants to study the essential roles of these closely related transcription factors. Using RT-PCR, whole-mount *in situ* hybridization (WISH), and fluorescence *in situ* hybridization, we examine *gsx1* and *gsx2* expression from early embryonic to late larval stages. *gsx1* is expressed initially in the hindbrain and diencephalon and later in the optic tectum, pretectum, and cerebellar plate. Comparatively, *gsx2* is expressed in the early telencephalon and later in the pallium and olfactory bulb. *gsx1* and *gsx2* are regionally co-expressed in the hypothalamus, preoptic area, and hindbrain, however rarely co-localize in the same cells. To identify forebrain target genes, we utilize mutants made with Transcription activator-like effector nucleases (TALEN). *gsx1* mutant zebrafish exhibit stunted growth, however, they survive through adulthood and are fertile. *gsx2* mutant zebrafish experience swim bladder inflation failure that prevents survival past larval stage. Using WISH and RT-qPCR we demonstrate altered expression of genes including, *distal-less homeobox* genes and *forkhead box* gene *foxp2*. This work provides novel tools with which other target genes and functions of Gsx1 and Gsx2 can be characterized across the CNS to better understand the unique and overlapping roles of these highly conserved transcription factors.

## INTRODUCTION

Central nervous system (CNS) development is a complex process wherein regionally expressed transcription factors contribute significantly in determining initial neuronal cell identity, connectivity, and function^1–3^. Transcription factors act coordinately to activate or repress target gene expression in progenitor cell domains^4,5^. Differential gene expression amongst neural progenitors generates distinct cell types and specifies neuronal properties, such as cell neurotransmitter content as seen in mouse^6–8^, chicken^9^, and zebrafish^10–12^. This process ultimately imparts initial identity to mature neuronal cells and forms the basis for neural circuit assembly and function. Thus, defining the spatiotemporal expression patterns and essential roles of vertebrate transcription factors is important for elucidating the functional mechanisms governing neurodevelopment. More importantly, these studies can provide fundamental insights about the molecular genetic contributions to the diverse neuroanatomical and behavioral phenotypes that are associated with neurodevelopmental disorders.

*genomic screen homeobox 1* and *2* (*gsx1* and *gsx2*, previously *gsh1* and *gsh2*) are closely related genes encoding homeobox transcription factors expressed in the CNS that were discovered in a screen for novel, non-clustered homeobox genes in mouse^13^. Homeobox genes characteristically encode transcription factors with a conserved 60-amino acid DNA-binding homeodomain^14,15^. Genes such as the *hox* genes specify cell types and body structures along the anterior-posterior (AP) axis in many species in patterns collinear with their 5’ to 3’ chromosomal positions within gene clusters^15–18^. As non-clustered and pseudo-clustered genes, *gsx1* and *gsx2* encode homeodomains with high (>80%) similarity to the *hox* genes^4,5,19^. *gsx1* and *gsx2* are the vertebrate homologs of *Drosophila melanogaster intermediate neuroblasts defective* (*ind*). *ind* and the *gsx* genes similarly regulate dorsoventral (DV) patterning^20–22^, and Ind and murine GSX2 elicit similar regulatory outcomes based on monomer versus homodimer DNA binding^23^. Interestingly, *ind* and the *gsx* genes are expressed in similar patterns in the fly neuroectoderm^20^, mouse neural tube^24^, and *Xenopus* neural plate^25^, supporting models for conserved neuroaxis domain specification across species.

Expression of *gsx1* and *gsx2* has been described in several vertebrates in varied detail. *gsx1* expression patterns are highly conserved across species, beginning in the hindbrain during somitogenesis in mouse^24^, *Xenopus*^25^, medaka^26^, and zebrafish^27^. During early embryonic stages in mouse *Gsx1* is expressed in the diencephalon and telencephalon and expands to the hypothalamus, thalamus, optic stalk, medulla, pons, and cerebellum^24^. Early expression in *Xenopus*, medaka^26^, and zebrafish^27,28^ occurs in similar regions such as the hypothalamus, olfactory bulb, optic tectum, and cerebellum. *gsx1* is also expressed as two dorsolateral stripes in the hindbrain and in the intermediate spinal cord in mice^24^, medaka^26^, and zebrafish^10^. *gsx2* is first detected slightly later than *gsx1* in the telencephalon and mesencephalon in mice and in the hindbrain in *Xenopus*^25,29^. Throughout neurodevelopment *gsx2* is expressed in the telencephalon, thalamus, hypothalamus, and cerebellum in mouse^29^, *Xenopus*^25^, and zebrafish^30^. Like *gsx1*, expression of *gsx2* appears similarly across species as two dorsolateral stripes in the hindbrain. *Gsx2* is expressed dorsal to *Gsx1* in the hindbrain in *Xenopus*^25^ and in the spinal cord of zebrafish^10^, consistent with their roles in DV patterning. Outside of the aforementioned reports^10,27,28,30^, expression of zebrafish *gsx1* and *gsx2* has not been comprehensively characterized and compared across all embryonic and early larval stages. Here we capitalize on the zebrafish model, which allows whole brain *in vivo* examination of expression to rigorously define the *gsx1* and *gsx2* expression profile. Defining a more complete expression profile of the *gsx* genes in zebrafish is an important step forward in elucidating critical Gsx1 and Gsx2 functions.

GSX1 and GSX2 promote regional neuronal identity in the ventral telencephalon and regulate the development of cortical, striatal, and olfactory bulb interneurons in mice^21,31–40^. Despite having similar roles in progenitor specification, GSX1 and GSX2 differentially regulate progenitor maturation; *Gsx2* maintains progenitors in an undifferentiated state while *Gsx1* promotes maturation by downregulating *Gsx2*^22,34^. *Gsx1* is implicated in hypothalamic and pituitary development, as knockout (KO) mice display a dwarf phenotype, reduced pituitary size, hormonal imbalances, and only survive a few weeks post-birth^41^. Consistently, *Gsx1* specifies multiple types of neuropeptidergic neurons in the arcuate nucleus of the hypothalamus^42^. *Gsx2* mouse KOs do not survive more than one day following birth, exhibit disturbed forebrain and hindbrain morphology^43^, and have expanded *Gsx1* expression in the ventral telencephalon^44^. Interestingly, *Gsx1* and *Gsx2* double KO mice display more severe forebrain phenotypes than *Gsx2* single KOs, supporting a model in which GSX1 partially compensates for loss of GSX2 function^21,44,45^. While much is known in the mouse forebrain, many key neurodevelopmental roles for *gsx1* and *gsx2* remain unknown, and roles for these transcription factors across the CNS have yet to be fully characterized in any vertebrate.

Outside of the forebrain, limited functional roles are reported for GSX1 and GSX2 in mammalian and non-mammalian model systems. *Gsx1* regulates an identity switch in mouse cerebellar neuronal progenitors in part through BMP/SMAD signaling^46,47^. Through a Notch signaling dependent mechanism, *Gsx1* and *Gsx2* regulate the temporal specification of glutamatergic and GABAergic interneurons in the mouse spinal cord^7^. In this region *Gsx1* also promotes neural stem and progenitor cell generation and decreases reactive glial scar formation to facilitate recovery from injury^48^. *gsx1* is implicated as a molecular marker of glutamatergic interneurons in the dorsal brainstem in zebrafish that regulate the acoustic startle response, and zebrafish with ablated *gsx1*-expressing neurons and mouse *Gsx1* knockouts similarly exhibit disrupted responsiveness to single and paired pulse acoustic-vibrational stimuli^28,49^. In zebrafish, *gsx2* is required for specification of neurons in the inferior olivary nuclei of the medulla^30^. *gsx1* and *gsx2* mark specific progenitor domains in the spinal cord of transgenic zebrafish similarly to mouse, with *gsx1* domains specifying glutamatergic, GABAergic, and glycinergic fates, and *gsx2* domains specifying glutamatergic fates only^10^.

Some GSX1 and GSX2 transcriptional target genes have been reported in the mouse forebrain and other brain regions^24,29,45,50,51^. However, target gene regulation by Gsx1 and Gsx2 across many brain regions has been understudied across vertebrates, including zebrafish. Several zebrafish orthologs for mouse GSX1 and GSX2 target genes exist, one example being *Distal-less homeobox 2* (*Dlx2*). Two paralogs, *dlx2a* and *dlx2b,* are found in the zebrafish genome, with *dlx2a* predicted to be the ortholog of mammalian *Dlx2*^52^. In mouse, GSX2 promotes *Dlx2* expression in the ventral telencephalon, while DLX2 in turn represses *Gsx1* and *Gsx2*^51^, and collectively this promotes ventral identity and mediates proliferative characteristics. Removal of *Gsx1* or *Gsx2* from a *Dlx1* and *Dlx2* double mutant background rescues some phenotypes observed, demonstrating that GSX/DLX inter-regulation is required for appropriate forebrain patterning. Two other *Dlx* genes, *Dlx5* and *Dlx6*, are similarly expressed in the forebrain of mice and zebrafish^52,53^. The *Dlx* genes coordinately regulate patterning of inhibitory neurons in the forebrain^52–54^, and importantly, the *DLX, FOX*, and other families of forebrain transcription factor encoding genes are implicated in aberrant neuronal signaling observed in patients with various neurodevelopmental disorders (NDDs)^1,40,55–58^. As such, it is important to investigate putative target genes for Gsx1 and Gsx2 to better understand their roles across brain regions during vertebrate neurodevelopment. In fact, the zebrafish model provides a tool with which this can be done rapidly and from the earliest neurodevelopmental time point possible.

In this study, we comprehensively resolve the neurodevelopmental expression of *gsx1* and *gsx2* in the zebrafish brain from early embryonic to late larval stages. Using *gsx1* and *gsx2* zebrafish mutants made using TALEN, we also demonstrate that *dlx2a, dlx2b, dlx5a,* and *dlx6a* are differentially regulated by Gsx1 and Gsx2. We further demonstrate that *forkhead box P2* (*foxp2*), a gene that is expressed in the mammalian and zebrafish CNS^59,60^ and is implicated in language deficits^58^, is regulated by Gsx2 in the zebrafish telencephalon. These studies are significant in that they establish novel tools for investigating Gsx1 and Gsx2 function during neurodevelopment and beyond in zebrafish across CNS regions.

## MATERIALS AND METHODS

### Zebrafish husbandry

All aspects of this study were approved by the West Virginia University IACUC. Adult zebrafish were maintained on a 14h/10h light/dark cycle at water temperature at 28-29°C. Breeding was performed using 1-liter breeding chambers with dividers (Aquaneering). Embryos were raised in 90×15mm petri dishes at 28.5°C in E3 media (pH 7.4; 0.005M NaCl, 0.00017M KCl, 0.00033M CaCl, 0.00033M MgSO4.7H_2_0, 1.5mM HEPES) in an incubator operating on a 14h/10h light/dark cycle. Staging of embryos was performed using standard procedures^61^. The following strain was used: TL (Tupfel long fin).

### Bioinformatics

Gene and protein sequences for all genes were obtained from the NCBI database (https://www.ncbi.nlm.nih.gov; see supplemental table S1 for accession numbers) and aligned using Clustal Omega (https://www.ebi.ac.uk/Tools/msa/clustalo/). Geneious was used to construct the rooted phylogenetic tree (https://www.geneious.com/academic/). The UCSF Genome Browser (http://genome.ucsc.edu/) and Ensembl database (http://uswest.ensembl.org/index.html) were used to evaluate exon and intron structures.

### Identification of zebrafish *gsx1* and *gsx2* mRNA transcripts using RT-PCR

Embryos and larvae obtained from TL crosses were raised to the desired ages (3.5 hpf-120 hpf), euthanized, frozen in liquid nitrogen, and stored at −80°C. Total RNA was extracted from 30 embryos and larvae at each age using a phenol chloroform extraction method with TRI-Reagent (Invitrogen). 1μg of total RNA was used with oligoDT to synthesize cDNA libraries (Superscript II First-Strand Synthesis kit, Invitrogen). 2μg of cDNA was used in 28 cycles of PCR with PlatinumTaq (Invitrogen) and intron-spanning gene-specific primers (see supplemental table S2). Amplicons were visualized and imaged using a FluorChemQ imager (ProteinSimple) on a 2% agarose gel with SYBR Safe DNA gel stain (Invitrogen) and excised using a blue light transilluminator (Clare Chemical Research). Sanger sequencing was used to confirm identity with NCBI sequences.

### Whole-mount *in situ* hybridization (WISH)

Embryos and larvae were raised to the desired ages and supplemented with 0.003% phenylthiourea (PTU) in E3 after 6 hpf to prevent pigmentation. For embryos younger than 48 hpf, chorions were removed by incubating in 50μg/mL Pronase (Sigma) at 28.5°C for 15 minutes. Embryos and larvae were anesthetized and fixed in cold 4% paraformaldehyde (PFA) overnight at 4°C. Fixed embryos were dehydrated using an increasing methanol wash series in 1xPBS (0%, 50%, 100% vol/vol methanol) and stored at −20°C in 100% MeOH for at least 24 hours and up to one year. The *gsx1*^27^ and *dlx2a*^62^ probes have been previously reported and were kind gifts of the Eisen and Karlstrom zebrafish labs. The probes for *gsx2*, *dlx2b*, *dlx5a*, *dlx6a*, and *foxp2* were designed in our lab. To generate antisense mRNA probes for *gsx2* and *dlx2b*, 1μg of age-specific total RNA was used with Invitrogen’s SuperScript III One-Step RT-PCR kit and gene-specific primers (see supplemental table S2) to amplify cDNA in 35 cycles of PCR. Amplicons were separated by 1% agarose gel electrophoresis, extracted using a QIAquick Gel Extraction kit (Qiagen), and subcloned into a pCR2.1_TOPO 4.0kb vector (Invitrogen). Sanger sequencing was used to confirm insert identity and directionality. 5μg of each plasmid was linearized with EcoR1 (*gsx2*) and Not1 (*dlx2b*) and probes were transcribed *in vitro* using SP6 polymerase (mMESSAGE mMACHINE kit, Ambion). To generate antisense mRNA probes for *dlx5a*, *dlx6a*, and *foxp2*, 1 μg of age-specific total RNA was used to synthesize cDNA libraries with oligoDT (Superscript II First-Strand Synthesis kit, Invitrogen). 1μg of cDNA was then used with gene-specific primers (see supplemental table S2) in 36 cycles of PCR. Amplicons were visualized using a 1% agarose gel with SYBR Safe DNA gel stain and purified using Qiagen’s QIAquick Gel Extraction kit. Probes were transcribed *in vitro* directly from purified amplicons using T7 polymerase (*dlx5a* and *dlx6a*) and SP6 polymerase (*foxp2*) (mMESSAGE mMACHINE kit, Ambion). The protocol for colorimetric WISH was adapted from Thisse and Thisse^63^ and performed essentially as in Bergeron *et al*.^28^. Embryos were hybridized with a digoxigenin (DIG)-tagged antisense mRNA probes detected by an anti-DIG antibody (Roche) and developed in NBT/BCIP (Roche). Staining was stopped by post-fixation in cold 4% PFA overnight at 4°C. Stained embryos were cleared in 75% glycerol and stored at 4°C protected from light.

### Fluorescence *in situ* hybridization (FISH)

FISH procedures were performed according to the *In Situ* Hybridization Chain Reaction v3.0 protocol (Molecular Instruments, Los Angeles, CA)^64^. Embryos were simultaneously hybridized with *gsx1* and *gsx2* probes (designed by Molecular Instruments) diluted in probe hybridization buffer overnight at 37°C. Excess probe was washed off the following day using probe wash buffer. Embryos were incubated for 30 minutes at room temperature in amplification buffer before adding the provided Alexa hairpins specific to the *gsx1* (Alexa Fluor 488) and *gsx2* (Alexa Fluor 546) mRNA sequences and incubating overnight at room temperature. Embryos were washed using 5x SSCT (5x SSC + 0.1% Tween 20) and stored in 5x SSCT at 4°C protected from light.

### Generation of *gsx1* and *gsx2* TALEN mutants

TALEN were designed using the freely available TALE-NT website that was created and is maintained by labs at Cornell University^65,66^. TALEN assembly was carried out using the Golden Gate vector system^66^ and separate destination vectors containing a modified FokI domain^67^. 100pg of *in vitro* transcribed mRNA (mMESSAGE mMACHINE kit, Ambion) was injected into the cell of each 1-cell stage zebrafish embryo to create G0. TALEN efficacy was checked by amplifying a 436bp fragment around the *gsx1* target site and a 409bp fragment around the *gsx2* target site using gene-specific primers (see supplemental table S2), followed by restriction digest of these amplicons using BtsI and EcoRI enzymes (NEB) respectively. Disruption of endonuclease cutting as evidenced by the presence of a full-length amplicon was considered effective, and siblings of these embryos were raised to adulthood and screened by crossing to wild-type (TL strain) adults to generate F1 offspring with single *gsx1* and *gsx2* mutant alleles. These alleles were sequence confirmed by DNA extraction from a subset of pooled sibling F1 embryos and PCR using gene-specific primers (see supplemental table S2), TOPO-TA sub cloning (Invitrogen), DH5α transformation, and Sanger sequencing of individual clones. F1 siblings carrying predicted loss of function mutations were raised to adulthood, genotyped, and crossed together to produce homozygous F2 for each new allele as a first pass mutant screen. Mutant lines are maintained by continuously crossing carriers to TL to eliminate possible off-target mutations over time, however, no off-target sites were predicted by TALE-NT.

### Genotyping for *gsx1Δ11* and *gsx2Δ13a* alleles

When genotyping was required to distinguish isolated *gsx1* and *gsx2* alleles, tissue was dissected from the most posterior end of the tail to use in DNA preparation. Tail tissue was denatured at 95°C for 10 minutes in DNA lysis buffer (10mM Tris pH 7.5, 50mM KCl, 0.3% Tween20, 0.3% Triton X, 1mM EDTA) and digested using 2mg/mL proteinase-K (Omega) at 55°C for at least 2h to overnight. Proteinase-K was heat-inactivated at 95°C for 10 minutes before the DNA was used in a standard DreamTaq (Thermo) PCR reaction with gene-specific primers (see supplemental table S2). Amplicons were visualized and imaged using a Syngene NuGenius imager with a blue light transilluminator (Clare Chemical Research) on a 4% agarose gel with SYBR Safe DNA gel stain. *gsx1Δ11* wild-type individuals have one band (140bp), mutants have one band (129bp), and heterozygotes have two bands at both sizes. *gsx2Δ13a* wild-type individuals have one band (134bp), mutants have one band (121bp), and heterozygotes have two bands at both sizes.

### *in silico* analyses and expression quantification

Sequences for *dlx2a*, *dlx2b*, *dlx5a*, *dlx6a*, and *foxp2* were identified in the NCBI database (see supplemental table S1 for accession numbers). The 25kb region upstream of the 5’ UTR of each gene was collected using Ensembl (https://useast.ensembl.org/index.html) and entered into ApE (http://jorgensen.biology.utah.edu/wayned/ape/). Assuming conservation of the GSX1^24^ and GSX2^29^ enhancer sequences identified in mouse, a 25kb region upstream of each gene 5’UTR was scanned for enhancer sequence variants. Annotated gene body schematics for the *Dlx2* orthologs were designed in Inkscape (https://inkscape.org/) and drawn using sequence information from Ensembl and ApE. Gene expression area was measured using FIJI-ImageJ by tracing the area of staining in the diencephalon or telencephalon. The telencephalon area was measured and used as a proxy for head size to correct for embryo size differences.

### Quantification of *dlx2a* and *dlx2b* expression using RT-qPCR

Embryos derived from heterozygous *gsx1Δ11* and/or *gsx2Δ13a* adults were euthanized and dissected at 30 hpf in cold RNAlater (Sigma) chilled by housing a 60×15mm petri dish on ice. A dissection anterior to the spinal cord was made to separate the head from the tail. Heads were stored in RNAlater at 4°C for up to one month and tails were used for DNA extraction and genotyping as previously described. 10-12 embryo heads of the same genotype were combined in a single 1.5mL snap tube and total RNA was extracted using a phenol chloroform extraction method with TRI-Reagent. 0.5μg of total RNA was used to synthesize cDNA libraries using oligoDT (Superscript II First-Strand Synthesis kit, Invitrogen). 1μL of cDNA was then used in a standard SYBR Green (Bio-Rad) qPCR reaction using gene-specific primers for *ef1a, dlx2a,* and *dlx2b* (see supplemental table S2). Samples were run on a Bio-Rad CFX Connect Real Time System using Bio-Rad CFX Maestro 1.1 Software. The 2^−ΔΔCt^ method^68^ was used to analyze raw Ct values and calculate gene expression changes relative to the housekeeping gene *ef1a*^69^.

### Microscopy and imaging

For WISH, embryos at 12 hpf were imaged at 6.3x on a Zeiss Stereo Discovery V.8 dissecting scope with an Axiocam 105 Color camera and analyzed using the ZEN 2.3 Lite software. Embryos of the remaining ages (24 hpf-144 hpf) were dissected and mounted in 75% glycerol under glass coverslips and imaged at 20x on a compound Zeiss Observer.Z1 with an Axiocam 503 Color camera. Imaging of genotyped samples for mutant studies was done blind by using a numeric code that could be aligned with genotype afterwards.

For FISH, embryos were dissected and mounted in 1x PBS under glass coverslips and imaged on an Olympus BX61 confocal microscope with Fluoview FV100 software. Imaging objectives were interchanged depending upon the area being investigated (Olympus UPlanApo, 20x or 40x oil immersion objectives with Olympus Immoil F30CC). Fluorophores used were Alexa Fluor 488 (*gsx1*) and Alexa Fluor 546 (*gsx2*).

For standard length measurements, embryos from heterozygous *gsx1Δ11* adults were raised and imaged at 4 dpf, 14 dpf, 1 month, 2 months, and 3 months old. Fish were anesthetized using MS-222, embedded in 1.5% low melt agarose in E3, and imaged next to a ruler on a Zeiss Stereo Discovery V.8 dissecting scope with an Axiocam 105 Color camera. FIJI-ImageJ was used to measure standard length^70^.

For swim bladder inflation studies, embryos derived from heterozygous *gsx1Δ11* or *gsx2Δ13a* crosses were raised under standard rearing conditions in 60×15mm petri dishes with 10mL of E3 media at a density of 30 embryos per dish. The number of larvae with and without inflated swim bladders were counted from days 3-6, and E3 was refreshed daily. Larvae were imaged on a Zeiss Stereo Discovery V.8 dissecting scope with an Axiocam 105 Color camera.

### Statistics

One-way ANOVAs with multiple comparisons and post hoc Tukey tests at α = 0.05 were performed in SPSS to evaluate significant differences between genotypes for the *gsx1* growth study and WISH expression analyses. For RT-qPCR studies, independent two-tailed t-tests at α = 0.05 were performed in SPSS to evaluate significant differences between 2^−ΔΔCt^ values calculated from raw Ct values. A Chi-square test (Pearson’s test) was performed using GraphPad to evaluate the association of genotype and swim bladder inflation at α = 0.05. Outliers for the growth study and WISH expression analyses were identified using GraphPad (Grubb’s test) and removed.

## RESULTS

### *gsx1* and *gsx2* expression in zebrafish embryos and larvae

To assess similarity between the Gsx1 and Gsx2 protein sequences in zebrafish, mouse, and human, we used a bioinformatics approach. We found that zebrafish Gsx1 shares 57/60 (95%) amino acids in the homeodomain with human and mouse GSX1 (Fig 1A). Zebrafish Gsx1 also shares 57/60 (93%) amino acids in the homeodomain with zebrafish, mouse, and human Gsx2. Interestingly, the homeodomain sequence is 100% identical between zebrafish, mouse, and human Gsx2. A rooted phylogenetic tree containing published Gsx1 and Gsx2 protein sequences reveals that zebrafish Gsx1 and Gsx2 cluster with their mammalian orthologs and also displays evolutionary divergence from *Drosophila* ortholog Ind (Fig 1B).

**Fig 1.**
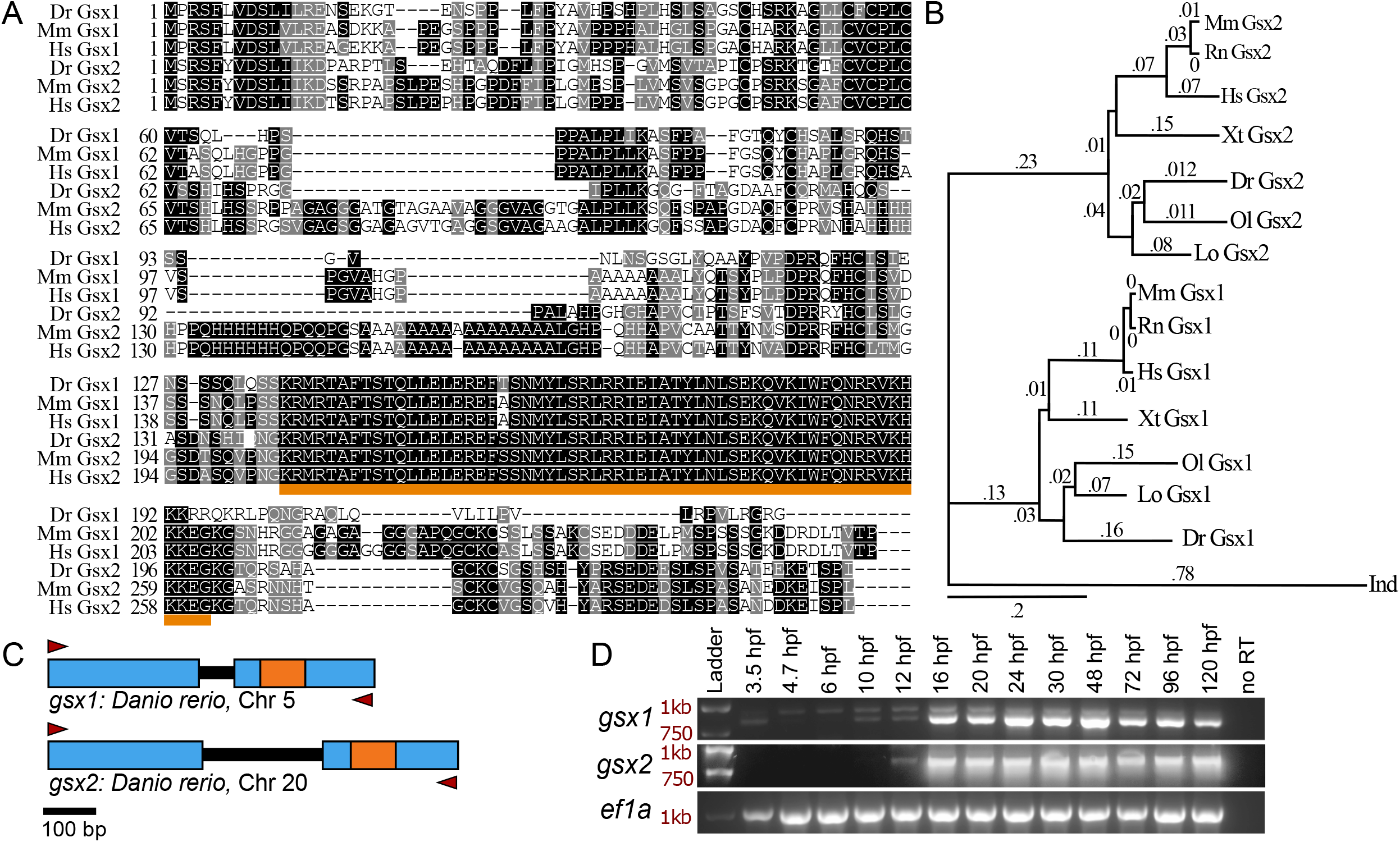
Gsx1 and Gsx2 are conserved across vertebrates and expressed in zebrafish from embryonic stages through adulthood. **A)** Amino acid sequence alignment of zebrafish, mouse, and human Gsx1 and Gsx2. Dr, *Danio rerio*; Mm, *Mus musculus*; Hs, *Homo sapiens*. Identical amino acids are shaded in black, similar amino acids are shaded in grey, and the region encoding the conserved DNA-binding homeodomain is underlined in orange. **B)** Rooted phylogenetic tree displaying the clustered relationship of published Gsx1 and Gsx2 protein sequences as well as a divergence from the Drosophila ortholog Ind. Values represent distance scores. Dm, *Drosophila melanogaster*; Lo, *Lepisosteus oculatus*; Ol, *Oryzias latipes*; Xt, *Xenopus tropicalis*. **C)** Schematic of the zebrafish *gsx1* and *gsx2* gene bodies. Blue boxes represent exons, black lines represent introns, and orange boxes represent the region encoding the homeodomain. Red arrowheads represent RT-PCR primer annealing sites. **D)** Agarose gels showing full-length cDNA transcripts of zebrafish *gsx1* (top), *gsx2* (middle), and *ef1a* (bottom) at specific ages. Red text indicates ladder sizes. Upper bands in the *gsx1* gel image represents trace genomic DNA that includes the 89 bp intron.

To document the neurodevelopmental time-course of *gsx1* and *gsx2* expression in zebrafish, we extracted total RNA from zebrafish embryos and larvae for use in RT-PCR (Fig 1C). *gsx1* expression was identified at 10 hours post fertilization (hpf), consistent with a previous report^27^, and persisted through our latest time point tested, 120 hpf (Fig 1D). Expression of zebrafish *gsx2* was first detected at 12 hpf and also persisted through 120 hpf. Interestingly, *gsx1* expression was observed at 3.5 hpf, suggestive of maternal contributions of *gsx1* to early embryonic development. However, analysis of maternal zygotic *gsx1* TALEN-generated mutants obtained through *in vitro* fertilization revealed that *gsx1* is not an essential maternal factor as maternal zygotic mutants are indistinguishable from zygotic mutants and develop identically (data provided by request). From these results, we can confirm that zebrafish *gsx1* and *gsx2* are expressed in zebrafish from embryonic to larval stages, suggesting an importance in early and later brain development and function.

### Expression of *gsx1* and *gsx2* during early development is complementary yet distinct

Known expression of *gsx1* in zebrafish is limited to select CNS regions beyond 30 hpf^10,27^ and expression of zebrafish *gsx2* is minimally reported from 48-72 hpf^30^. Outside of a transgenic analysis documenting *gsx1* and *gsx2* expression together in the 36-48 hpf spinal cord^10^, expression of *gsx1* and *gsx2* during neurodevelopment in zebrafish has not been comprehensively analyzed. We first used whole-mount *in situ* hybridization (WISH) to characterize and compare *gsx1* and *gsx2* expression in zebrafish embryos and larvae. Consistent with RT-PCR results, *gsx2* expression was detected at 12 hpf in the presumptive forebrain in the anterior neural plate (Fig 2H). At 12 hpf *gsx1* is expressed in the presumptive hindbrain in rhombomere 3 (Fig 2A), consistent with a previous report^27^. From 16-24 hpf *gsx2* expression is present in the diencephalon and telencephalon (Fig 2I-J), with 24 hpf marking the first appearance of *gsx2* expression in the caudal hindbrain (Fig 2K). Expression of *gsx2* in the zebrafish spinal cord is seen clearly in transgenic reporter lines at this time^10^, but we predict endogenous *gsx2* expression in this region is highly transient and difficult to detect by WISH. From 16-24 hpf *gsx1* expression is observed in the forebrain, midbrain, hindbrain, and spinal cord (Fig 2B-C). Conversely, *gsx1* is expressed across the rostral to caudal extent of the hindbrain at 24 hpf (Fig 2D). *gsx2* expression persists in the diencephalon, telencephalon, hindbrain, and spinal cord through 30 hpf (Fig 2L-M), and at this age *gsx1* is expressed in the diencephalon, midbrain, hindbrain, and spinal cord (Fig 2E-F). Dorsal views at this age reveal that *gsx1* and *gsx2* exist in two dorsolateral columns in the hindbrain (Fig 2O and S).

**Fig 2.**
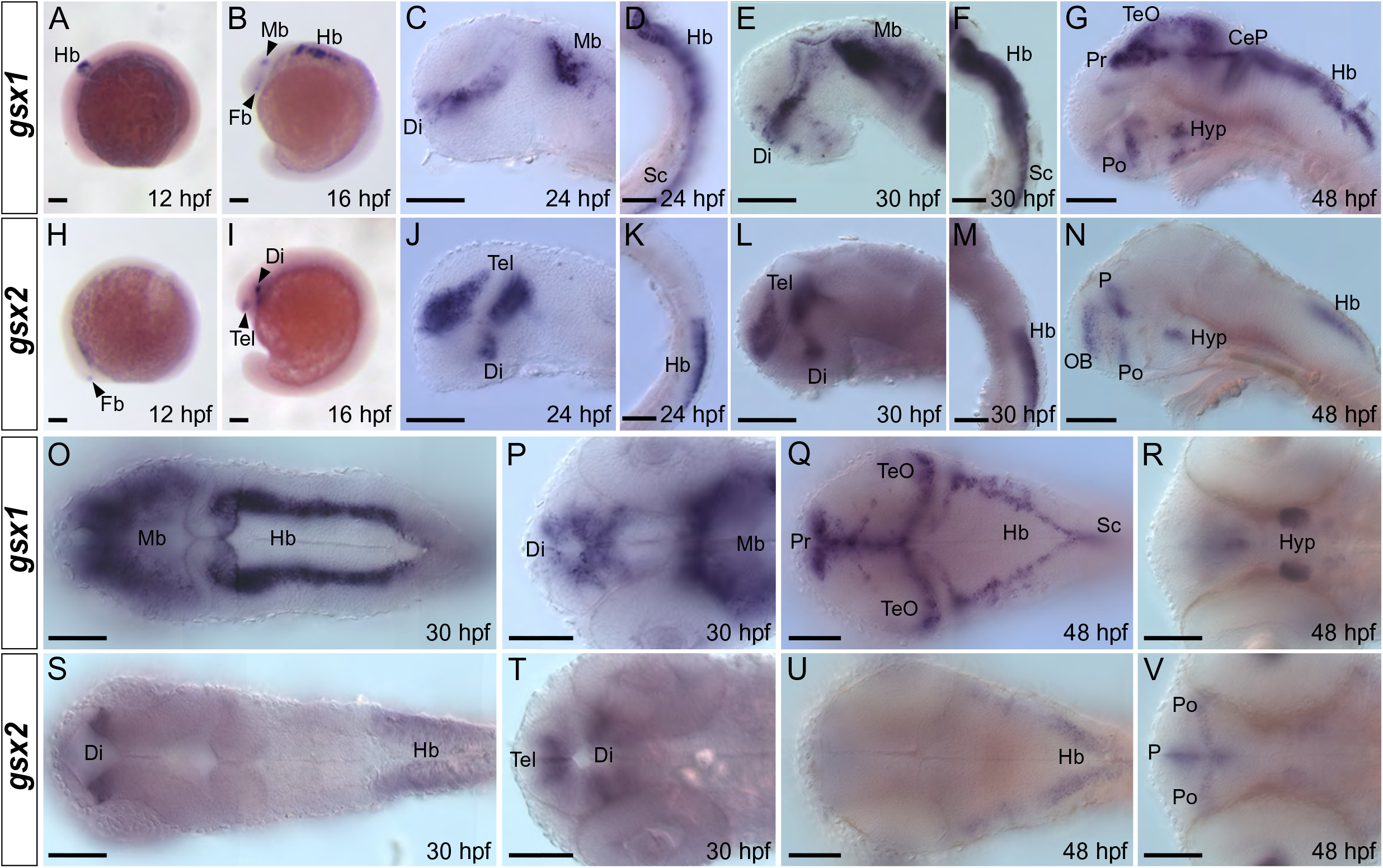
Expression of *gsx1* and *gsx2* in embryonic zebrafish is dynamic and unique. **A-G)** Lateral mounts showing expression of *gsx1* from 12-48 hpf. **H-N)** Lateral mounts showing expression of *gsx2* from 12-48 hpf. **O-R)** Dorsal (O & Q) and ventral (P & R) mounts showing expression of *gsx1* from 30-48 hpf. **S-V)** Dorsal (S & U) and ventral (T & V) mounts showing expression of *gsx2* from 30-48 hpf. A, B, H, and I are dissecting scope images and scale bar represents 500μm. Remaining images are compound scope images taken at 20X with samples mounted under cover glass and anterior facing left, eyes removed in lateral views. Scale bars represent 100μm. CeP = cerebellar plate, Di = diencephalon, Fb = forebrain, Hb = hindbrain, Hyp = hypothalamus, Mb = midbrain, OB = olfactory bulb, P = pallium, Po = preoptic area, Pr = pretectum, Sc = spinal cord, Tel = telencephalon, TeO = optic tectum.

Our observed *gsx1* expression through 30 hpf confirms previous findings^27^, however we continued characterizing *gsx1* expression through late embryonic and larval development along with *gsx2*. By 48 hpf, *gsx2* expression is restricted to the olfactory bulb, preoptic area, hypothalamus, pallium, and hindbrain (Fig 2N, 2U-V). At this age, *gsx1* expression is seen in the preoptic region, hypothalamus, pretectum, optic tectum, cerebellar plate, hindbrain, and spinal cord (Fig 2G, 2Q-R). At 72 hpf *gsx2* expression is faintly present in the pallium and olfactory bulb (Fig 3J-K), while *gsx1* expression persists in the pretectum, optic tectum, hypothalamus, and hindbrain (Fig 3A-C). Expression of *gsx2* through 4-5 days post fertilization (dpf) persists faintly in the pallium and hindbrain (Fig 3L-O), however *gsx1* expression strongly persists in the pretectum, optic tectum, hypothalamus, and hindbrain (Fig 3D-I). Collectively, these WISH analyses provide insight to the dynamic expression patterns of *gsx1* and *gsx2* during embryonic and larval stages in zebrafish.

**Fig 3.**
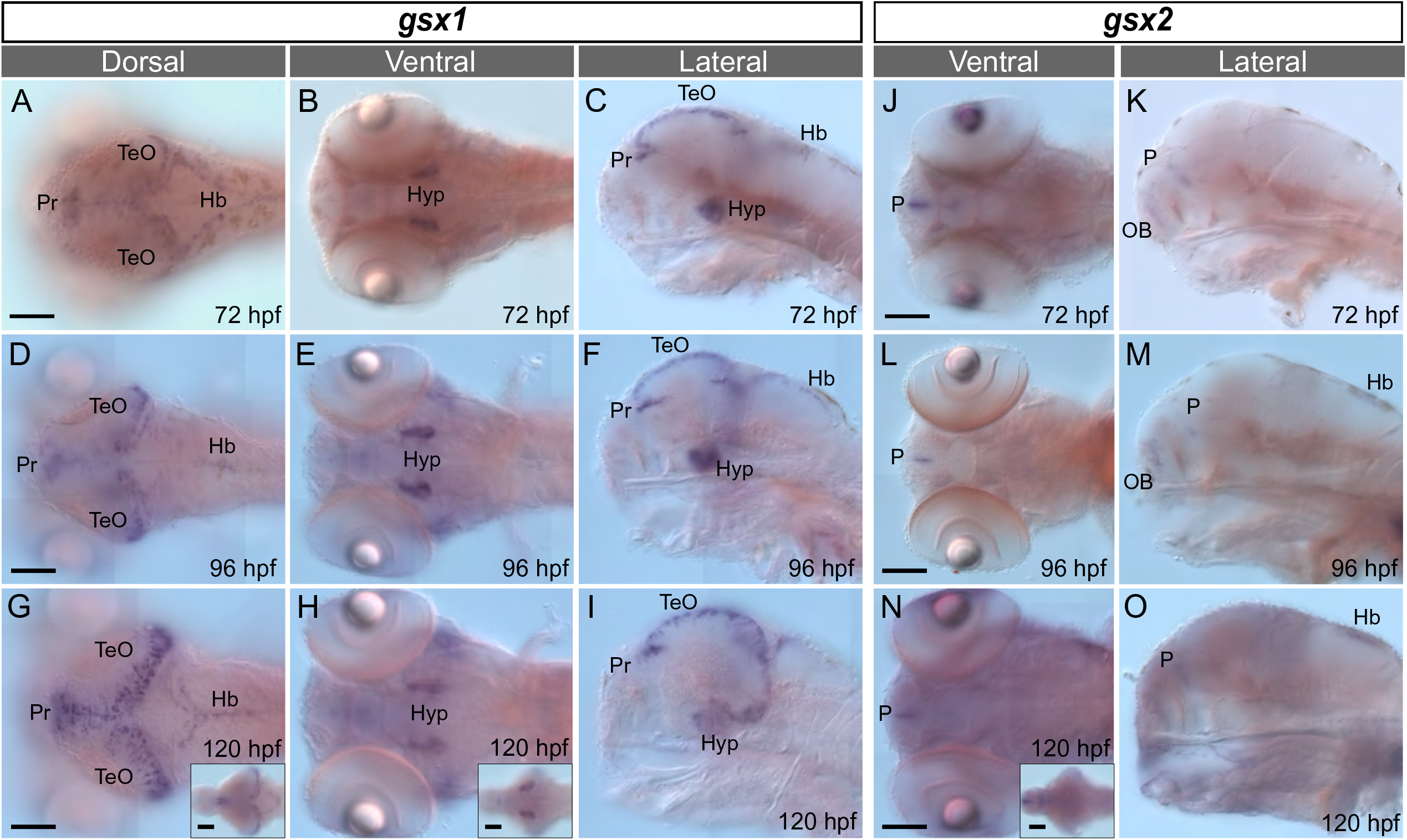
Expression of *gsx1* and *gsx2* is restricted in late embryonic and early larval stages. **A-I)** Expression of *gsx1* from 72-120 hpf. Leftmost column are dorsal views, middle column are ventral views, and rightmost column are lateral views. **J-O)** Expression of *gsx2* from 72-120 hpf. Left column are dorsal views and rightmost column are lateral views. All images are compound scope images taken at 20X with samples mounted under cover glass and anterior facing left, eyes removed in lateral views. Scale bars represent 100μm. Insets are whole brain dissections at the same age mounted dorsally. Hb = hindbrain, Hyp = hypothalamus, OB = olfactory bulb, P = pallium, Pr = pretectum, TeO = optic tectum.

### Co-localization of *gsx1* and *gsx2* is minimal

To be more precise in assessing co-localization of *gsx1* and *gsx2* in cells, we turned to fluorescence *in situ* hybridization (FISH) at embryonic and larval stages. At 24 hpf *gsx1* and *gsx2* are regionally co-expressed at the border of the dorsal diencephalon and ventral telencephalon (Fig 4A^i-iii^ and 4B^i-iii^; max z-projections), however they very minimally co-localize in the same cells (insets in Fig 4A^iii^ and 4B^iii^; single z-stack plane). At this age *gsx1* and *gsx2* are also regionally co-expressed in the hindbrain, with *gsx2* expressed dorsal to *gsx1* (Fig 4A^iv-vi^, 4B^iv-vi^). At 30 hpf, *gsx1* and *gsx2* are regionally co-expressed in the ventral diencephalon (Fig 4C^i-iii^ and 4D^i-iii^; max z-projections), however rarely co-localize in the same cells (insets in Fig 4C^iii^ and 4D^iii^; single z-stack plane). In the hindbrain at this age *gsx1* and *gsx2* remain segregated dorsoventrally (Figs 4C^iv-vi^ and 4D^iv-vi^; max z-projections) and rarely co-localize in the same cells (inset in Fig 4D^vi^; single z-stack plane).

**Fig 4.**
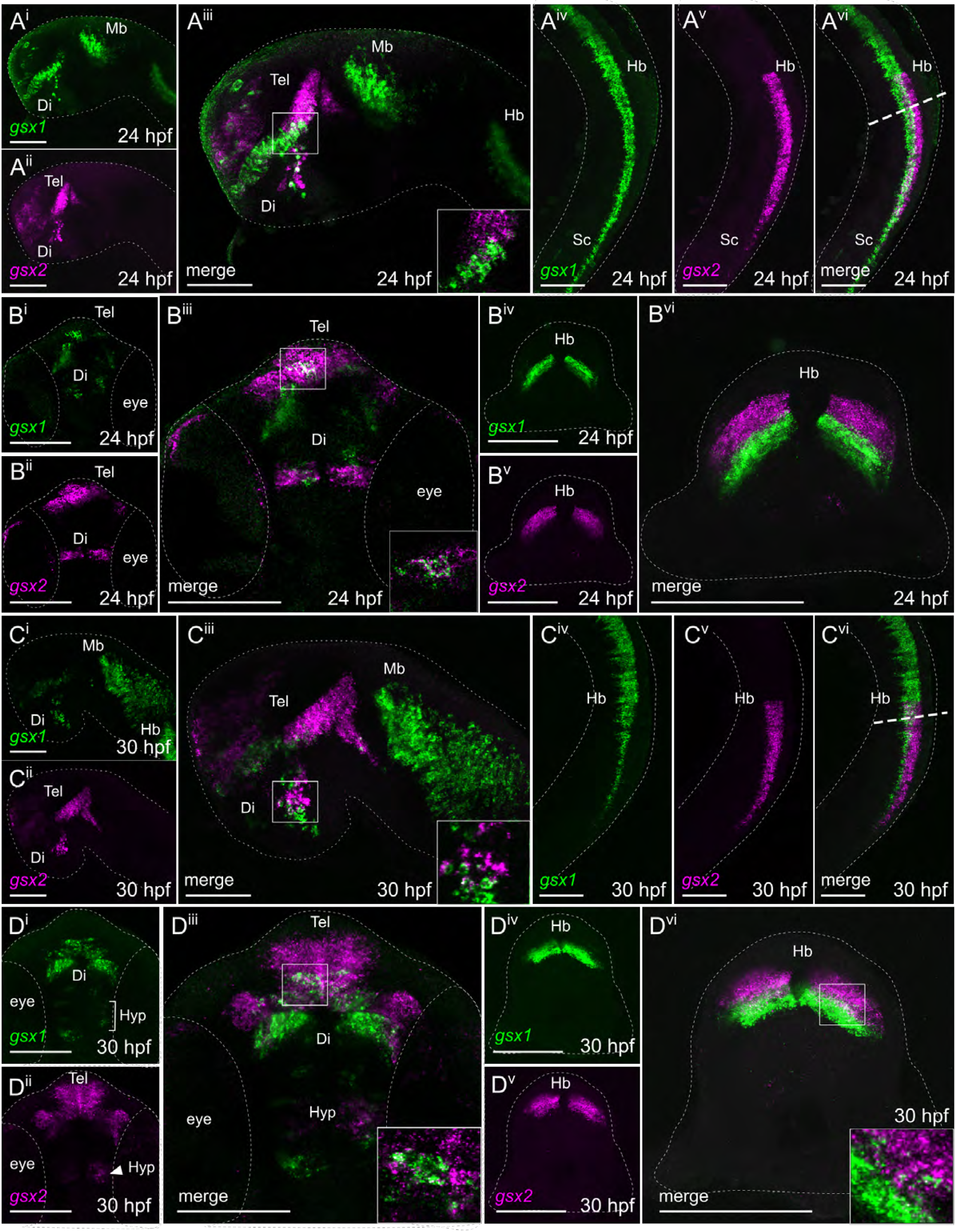
Fluorescence *in situ* hybridization confirms minimal co-localization of *gsx1* and *gsx2* during embryonic development. **A^i^-A^vi^**) Lateral views of *gsx1* and *gsx2* expression at 24 hpf. **B^i^-B^iii^)** Ventral view of *gsx1* and *gsx2* expression at 24 hpf. **B^iv^-B^vi^)** Cross section view taken at the dashed line in A^vi^. **C^i^-C^vi^)** Lateral views of *gsx1* and *gsx2* expression at 30 hpf. **D^i^-D^iii^)** Ventral view of *gsx1* and *gsx2* expression at 30 hpf. **D^iv^-D^vi^)** Cross section view taken at the dashed line in C^vi^. Lateral views were taken at 20X and ventral/cross section views were taken at 40X, all with anterior facing left. All were pseudocolored using FIJI ImageJ and scale bars represent 100μm. For lateral views, eyes were dissected off. Main images are max z-projections and insets are single z-stack slices zoomed into the boxed region shown in the main image. Di = diencephalon, Hb = hindbrain, Mb = midbrain, Sc = spinal cord, Tel = telencephalon.

By 48 hpf *gsx1* and *gsx2* are regionally co-expressed in the hypothalamus and preoptic area (Fig 5A^i-iii^ and 5B^i-iii^; max z-projections), however rarely co-localize in the same cells (insets in Fig 5A^iii^ and 5B^iii^; single z-stack plane). Distinct segregation of *gsx1* ventrally and *gsx2* dorsally in the hindbrain is still apparent at 48 hpf (Fig 5A^iv-vi^ and 5B^iv-vi^; max z-projections) however they rarely co-localize in the same cells (inset in Fig 5A^vi^; single z-stack plane). Interestingly, by this age *gsx1* expression appears to extend ventrally while *gsx2* expression remains isolated dorsally (Fig 5A^iv-vi^, 5B^iv-vi^). This finding is reminiscent of reported roles for *Gsx2* and *Gsx1* to regulate neuronal progenitor proliferation versus differentiation, respectively^22^, and we believe this ventral extension represents the outgrowth of projections from maturing neuronal progenitors. At 72 hpf regional co-expression of *gsx1* and *gsx2* is restricted to the preoptic area (Fig 5C^i-vi^; max z-projections), however again, co-localization in the same cells is minimal (Fig 5C^iii^; single z-stack plane).

**Fig 5.**
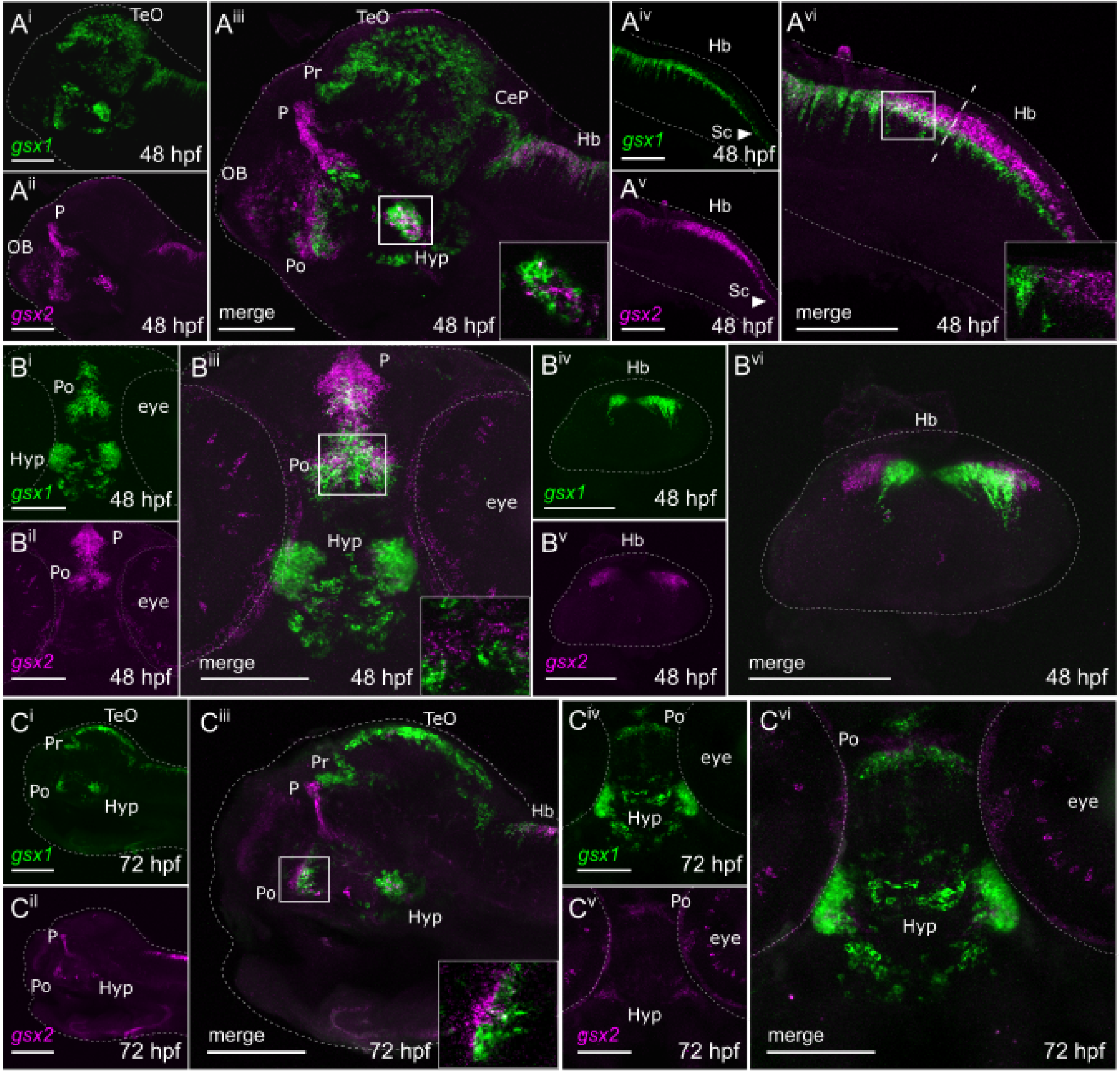
Fluorescence *in situ* hybridization confirms minimal co-localization of *gsx1* and *gsx2* during late embryonic and early larval development. **A^i^-A^vi^)** Lateral views showing expression of *gsx1* and *gsx2* at 48 hpf. **B^i^-B^ii^)** Ventral views showing expression of *gsx1* and *gsx2* at 48 hpf. **B^iv^-B^vi^)** Cross section view taken at the dashed line in A^vi^. **C^i^-C^iii^)** Lateral view showing *gsx1* and *gsx2* expression at 72 hpf. **C^iv^-C^vi^)** Ventral view showing *gsx1* and *gsx2* expression at 72 hpf. Lateral views were taken at 20X and ventral/cross section views were taken at 40X, all with anterior facing left. All were pseudocolored using FIJI ImageJ and scale bars represent 100μm. For lateral views, eyes were dissected off. Main images are max z-projections and insets are single z-stack slices zoomed into the boxed region shown in the main image. CeP = cerebellar plate, Di = diencephalon, Hb = hindbrain, Hyp = hypothalamus, Mb = midbrain, OB = olfactory bulb, P = pallium, Po = preoptic area, Pr = pretectum, Sc = spinal cord, Tel = telencephalon, TeO = optic tectum.

In brains dissected from 6 dpf larvae, we observed *gsx1* and *gsx2* expression patterns that were not directly apparent through colorimetric WISH analyses. *gsx1* expression appears in regions reminiscent of our later stage WISH analyses including the pretectum, hypothalamus, optic tectum, preoptic area, cerebellar plate, and hindbrain (Fig 6A-F, max z-projections). Dorsal views of the brain confirmed that *gsx2* is expressed in the pallium at this age (Fig 6A-C), and also revealed distinct expression in the hindbrain not clearly observed through WISH. Additionally, ventral views of the brain at this age also revealed that *gsx2* is regionally co-expressed with *gsx1* in the hypothalamus (Fig 6D-F) however they rarely co-localize in the same cells (Fig 6F; single z-stack plane). Combined, our WISH analyses reveal that *gsx1* and *gsx2* expression is dynamic throughout neurodevelopment, and FISH demonstrates for the first time that they largely exist in distinct cellular populations.

**Fig 6.**
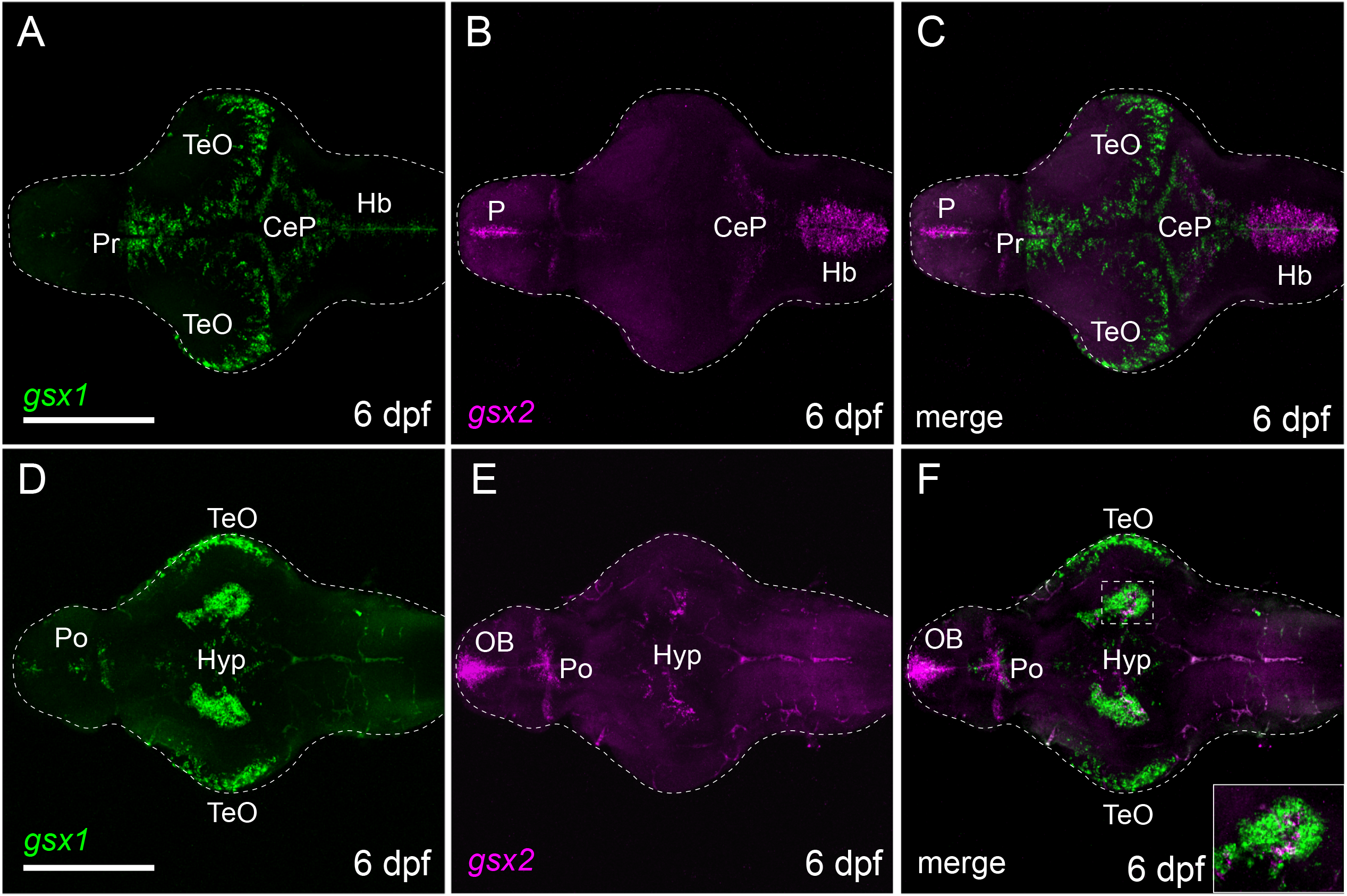
*gsx1* and *gsx2* are expressed through 6dpf in the brain. **A-C)** Dorsal views showing expression of *gsx1* and *gsx2* at 6 dpf. **D-F)** Ventral views showing expression of *gsx1* and *gsx2* at 6 dpf. Images taken at 20X with anterior facing left. All were pseudocolored using FIJI ImageJ and scale bars represent 100μm. Main images are max z-projections and insets are single z-stack slices zoomed into the boxed region shown in the main image. Hb = hindbrain, Hyp = hypothalamus, OB = olfactory bulb, P = pallium, Po = preoptic area, Pr = pretectum, Tel = telencephalon, TeO = optic tectum.

### *gsx1* and *gsx2* TALEN mutants exhibit unique phenotypes

There is limited knowledge about how Gsx1 and Gsx2 function across several developing brain regions where they are expressed in vertebrates and that we report by WISH and FISH in zebrafish. In mouse, loss of *Gsx1* leads to abnormal hypothalamic-pituitary signaling^41^, and mutations in *Gsx2* leads to disturbed forebrain morphology^43^. To further examine the roles of *gsx1* and *gsx2* in neurodevelopment, we generated zebrafish mutants using TALEN (Fig 7A). For *gsx1*, we generated alleles with an 11 base-pair (bp) deletion (*gsx1Δ11*), 5bp deletion (*gsx1Δ5*), and 1bp insertion (*gsx1Δpl1*). For *gsx2*, we generated alleles with a 13bp deletion (*gsx2Δ13a* and *gsx1Δ13b*) and 5bp deletion (*gsx2Δ5*). All mutations occur in the first exon of the zebrafish *gsx1* and *gsx2* genes and should result in premature stop codons and immature transcripts lacking the homeobox DNA binding domain. Phenotypes observed are thus far consistent with a loss of function across all *gsx1* and *gsx2* alleles.

**Fig 7.**
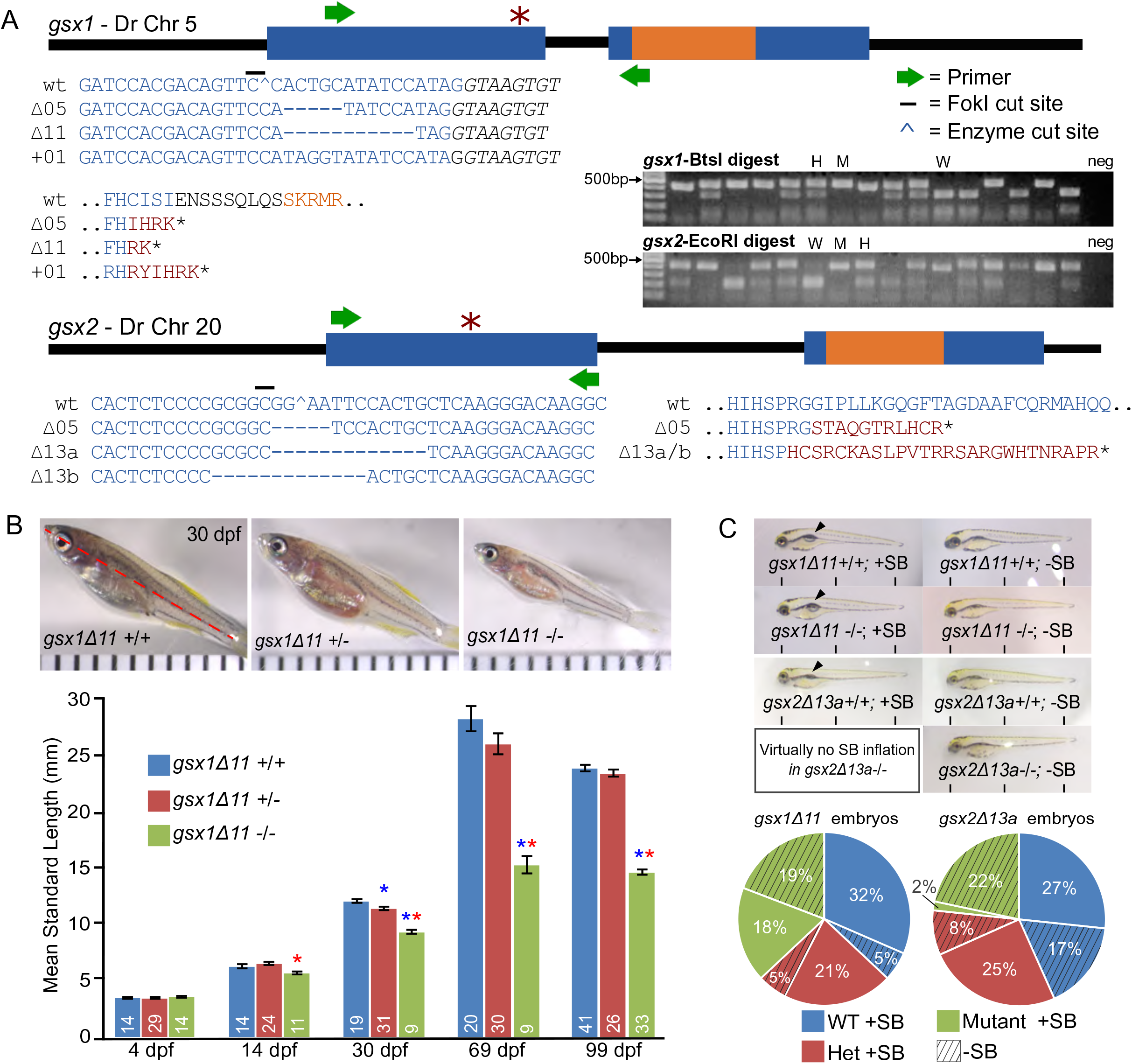
*gsx1* TALEN mutants experience stunted growth and *gsx2* TALEN mutants experience swim bladder inflation failure. **A)** Schematic of the *gsx1* and *gsx2* gene bodies and targeted TALEN mutation site (*). All mutations should result in a premature stop codon. Text color corresponds with gene body structure (blue = exon, black = intron, orange = homeodomain, red = mutant sequence). Inset gel shows restriction digest of *gsx1* and *gsx2* amplicons in wild-type (W), heterozygous (H) and mutant (M) individuals; endonuclease cutting is disrupted in mutants. **B)** Comparison of the standard length (SL, red line) of *gsx1Δ11* wild-type, heterozygous, and mutant siblings at 30 dpf. Images are dissecting scope images and distances between tick marks represent 1mm. Graph shows the quantification of the long-term growth study data from 4 dpf to 99 dpf. Asterisks of each color indicate significant differences between that group and the genotype of the same color (blue, wild-type; red, heterozygous; green, mutant). **C)** Top, comparison of *gsx1Δ11* and *gsx2Δ13a* wild-type and mutant larvae with (+SB) and without (-SB) swim bladders, respectively; Bottom, quantification of the percentage of larvae of each genotype with and without swim bladders (*gsx1* n=73, *gsx2* n=60).

Through assessing our *gsx1* mutant zebrafish, we found that these fish experience stunted growth starting at 14 dpf. No significant differences in standard length were found across genotypes in 4 dpf larvae (Fig 7B), however by 14 dpf standard length of *gsx1Δ11*−/− larvae was significantly smaller than *gsx1Δ11+/−* siblings (p=0.002). By one month *gsx1Δ11*−/− larvae were significantly smaller than both *gsx1Δ11*+/+ and *gsx1Δ11*+/− siblings (p<0.001 for both), and this difference persisted through 2 months (p<0.001 for both) and 3 months of age (p<0.001 for both). These analyses reveal a growth-related phenotype in *gsx1Δ11−/−* zebrafish similar to reports in mouse^41^. However, unlike *Gsx1* mutant mice, our *gsx1* mutant zebrafish survive to adulthood, allowing investigations of early and later Gsx1 function across brain regions.

Embryos derived from heterozygous *gsx2Δ13a* parents are indistinct from *gsx1Δ11+/−* cross embryos (Fig 7C, top). However, *gsx2Δ13a*−/− embryos largely fail to inflate their swim bladders by 6 dpf under standard rearing conditions, preventing their survival. There was a significant association between swim bladder inflation and genotype in offspring from *gsx2Δ13a+/−* crosses, as less *gsx2Δ13a*−/− larvae had inflated swim bladders compared to *gsx2Δ13a*+/+ and *gsx2Δ13a*+/− larvae (Fig 7C, bottom; X^2^=22.8, p<0.001). Swim bladder inflation did not differ between genotypes in *gsx1Δ11+/−* crosses (X^2^=.32, p=.851). These results demonstrate that swim bladder inflation failure is a result of a mutation in *gsx2,* and supports the important developmental role for Gsx2 in vertebrates, including zebrafish.

### Gsx1 and Gsx2 differentially regulate *distal-less homeobox 2a* and *2b*

Enhancer sequences have been reported for murine GSX1^24^ and GSX2^29^, and previous studies report *Distal-less homeobox 2* (*Dlx2*) as a target gene of GSX1 and GSX2 in the mouse forebrain^51^. *Dlx2* expression overlaps with *Gsx1* and *Gsx2* in the medial, caudal, and lateral ganglionic eminences (MGE, CGE, and LGE, respectively) of the mouse telencephalon where they coordinately regulate early neuronal progenitor patterning^40,71,72^. This work shows that GSX1 and GSX2 upregulate *Dlx2*^51^. Therefore, we sought to determine if the zebrafish ortholog *dlx2a* or its paralog *dlx2b* are Gsx1 and Gsx2 target genes. Published gene sequences for human, mouse, and zebrafish *Dlx2* were analyzed *in silico* for Gsx1 and Gsx2 enhancer sequences, which we assume are conserved in zebrafish. We found that human *DLX2*, mouse *Dlx2*, and zebrafish *dlx2b* possess putative Gsx1 and Gsx2 enhancer sequences upstream of their 5’UTRs (Fig 8A). Human *DLX2* and zebrafish *dlx2b* possess a putative enhancer sequence that both Gsx1 and Gsx2 could bind. Zebrafish *dlx2a* possesses putative Gsx2 enhancer sequences only.

**Fig 8.**
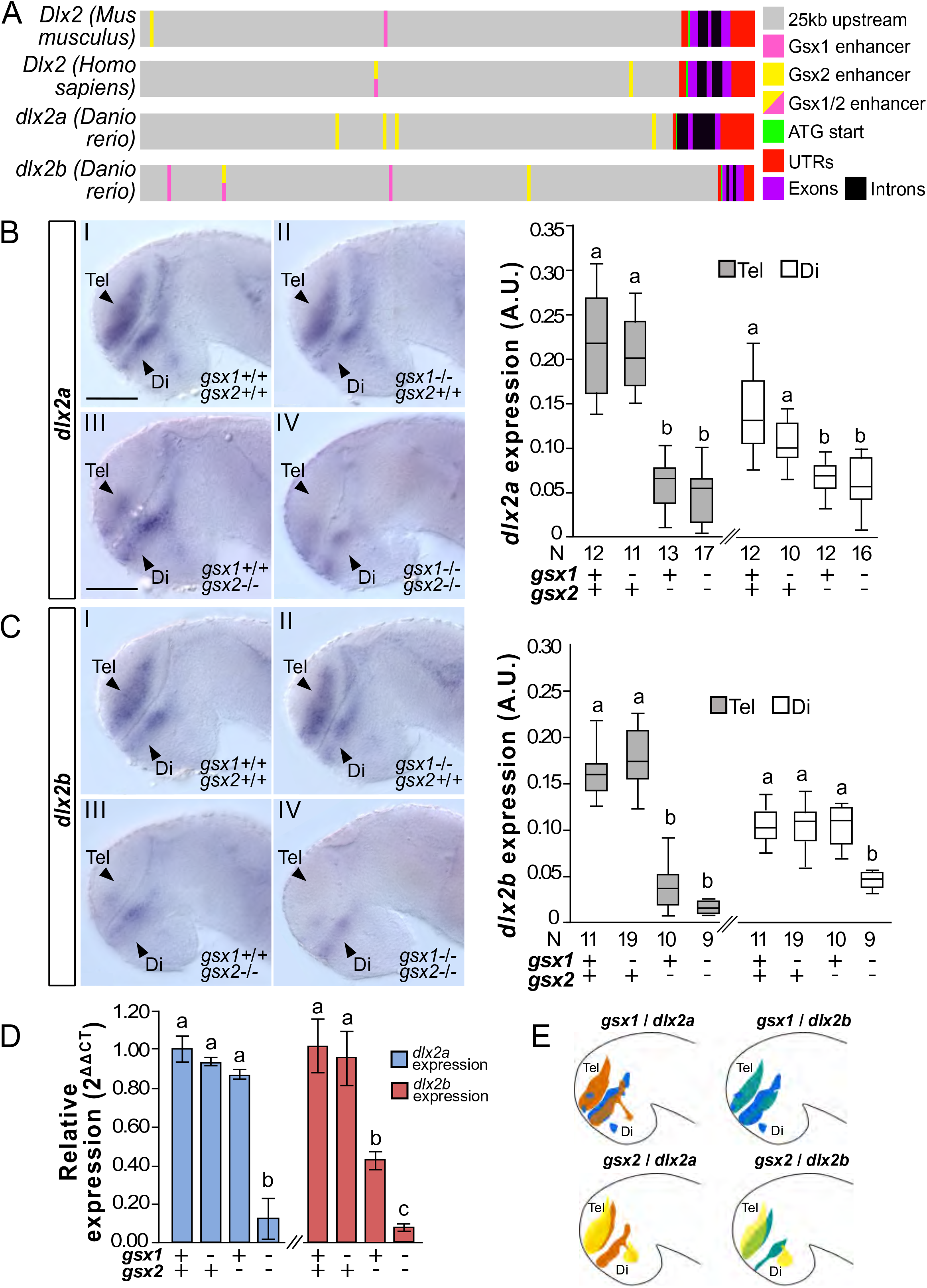
Gsx1 and Gsx2 differentially regulate *dlx2a* and *dlx2b* expression. **A)** Schematic of Gsx1 and 2 enhancer sequences in the Dlx2 orthologs. **B)** Left, *dlx2a* expression at 30 hpf in wild-type, *gsx1Δ11*−/−, *gsx2Δ13a*−/−, and *gsx1Δ11*−/−;*gsx2Δ13a*−/− zebrafish. Images are 20X compound scope images with samples mounted under cover glass, eyes dissected, and anterior facing left. Scale bar = 50μm. Right, FIJI-ImageJ quantification of *dlx2a* expression in the telencephalon (grey bars) and diencephalon (white bars). Genotypes and sample size are listed under the X axis. Different letters represent significant differences. **C)** Left, *dlx2b* expression at 30 hpf in wild-type, *gsx1Δ11*−/−, *gsx2Δ13a*−/−, and *gsx1Δ11*−/−/*gsx2Δ13a*−/− zebrafish; Right, quantification of *dlx2b* expression. **D)** RT-qPCR data showing relative expression of *dlx2a* (blue bars) and *dlx2b* (red bars) in wild-type, *gsx1Δ11*−/−, *gsx2Δ13a*−/−, and *gsx1Δ11*−/−;*gsx2Δ13a*−/− zebrafish compared to the reference gene *ef1a*. Different letters indicate significant differences within each target gene. **E)** Schematics of *gsx1*, *gsx2*, *dlx2a*, and *dlx2b* expression at 30 hpf.

To determine if Gsx1 or Gsx2 regulate *dlx2a* and/or *dlx2b* in zebrafish, we quantified *dlx2a* and *dlx2b* gene expression area in 30 hpf embryos yielded from *gsx1Δ11*+/−;*gsx2Δ13a*+/− crosses using WISH and RT-qPCR. We found that *dlx2a* expression is not significantly different between *gsx1Δ11*+/+;*gsx2Δ13a*+/+ and *gsx1Δ11*−/−;*gsx2Δ13a*+/+ embryos (Fig 8B^I-II^), however is significantly reduced in both *gsx1Δ11*+/+;*gsx2Δ13*−/− and *gsx1Δ11*−/−;*gsx2Δ13a*−/− embryos in the diencephalon (p<0.001 for both) and telencephalon (p<0.001 for both; Fig 8B^III-IV^ and graphs). Expression of *dlx2b* is not different between *gsx1Δ11*+/+;*gsx2Δ13a*+/+ and *gsx1Δ11*−/−;*gsx2Δ13a+/+* embryos (Fig 8B^I-II^), however is significantly reduced in both *gsx1Δ11*+/+;*gsx2Δ13*−/− and *gsx1Δ11*−/−;*gsx2Δ13a*−/− embryos in the diencephalon (p<0.001 for both) and telencephalon (p<0.001 for both; Fig 8B^III-IV^ and graphs).

Consistent with WISH, RT-qPCR revealed that *dlx2b* expression is significantly reduced in *gsx1Δ11*+/+;*gsx2Δ13a*−/− and *gsx1Δ11*−/−;*gsx2Δ13a*−/− embryos compared to wild-types (p=0.005 and 0.002, respectively; Fig 8D). Furthermore, we also observed that *dlx2b* expression is significantly reduced in *gsx1Δ11*−/−;*gsx2Δ13a*−/− embryos compared to *gsx1Δ11*+/+;*gsx2Δ13a*−/− embryos (p=0.012), suggesting that Gsx1 partially sustains *dlx2b* expression which becomes further reduced upon loss of both *gsx1* and *gsx2*. Unlike WISH, RT-qPCR showed that *dlx2a* expression is only significantly reduced in *gsx1Δ11*−/−;*gsx2Δ13a*−/− embryos (p=<0.001) and not in *gsx1Δ11*+/+;*gsx2Δ13a*−/− embryos (p=0.225; Fig 8D). WISH shows that in *gsx1Δ11*+/+;*gsx2Δ13a*−/− embryos, *dlx2a* and *dlx2b* expression is lost in the telencephalon, where *gsx2* is expressed, yet sustained in the diencephalon, where *gsx1* is expressed. This suggests that zebrafish Gsx1 and Gsx2 differentially regulate *dlx2a* and *dlx2b* expression in the telencephalon and diencephalon and that visible changes in *dlx2a* expression cannot be detected by RT-qPCR of whole brains at 30 hpf.

### Gsx1 and Gsx2 differentially regulate *dlx5a*, *dlx6a,* and *foxp2*

To identify additional Gsx1 and Gsx2 target genes in the zebrafish forebrain, we applied our *in silico* and WISH approaches to assess regulation of *dlx5a* and *dlx6a*, which are closely related to *dlx2a* and *dlx2b*. Both *dlx5a* and *dlx6a* are expressed in overlapping patterns with *dlx2a* and *dlx2b* in the zebrafish forebrain^52,57^ and coordinately regulate inhibitory neuron patterning in subpallial regions with the other *dlx* orthologs^73^. Published gene sequences for zebrafish *dlx5a* and *dlx6a* were analyzed *in silico* for putative Gsx1 and Gsx2 enhancer sequences. Zebrafish *dlx5a* possesses both Gsx1 and Gsx2 enhancer sequences in the 25kb region upstream of the 5’UTR, while zebrafish *dlx6a* only possesses Gsx2 enhancers (Fig S1A).

To determine if Gsx1 or Gsx2 regulate *dlx5a* and *dlx6a* in the zebrafish forebrain, we again quantified gene expression area in 30 hpf embryos yielded from *gsx1Δ11*+/−;*gsx2Δ13a*+/− crosses. We found that *dlx5a* is significantly reduced in both *gsx1Δ11*+/+;*gsx2Δ13*−/− and *gsx1Δ11*−/−;*gsx2Δ13a*−/− embryos in the telencephalon (p=<0.001), however only significantly reduced in *gsx1Δ11*−/−;*gsx2Δ13a*−/− embryos in the diencephalon (p=<0.001, Fig S1B^i-iv^, S1C). These results suggest that in the telencephalon, *dlx5a* is regulated by Gsx2, however in the diencephalon both Gsx1 and Gsx2 are required for normal expression. For *dlx6a*, we observed significant reductions in expression in in both *gsx1Δ11*+/+;*gsx2Δ13*−/− and *gsx1Δ11*−/−;*gsx2Δ13a*−/− embryos in the telencephalon only (p=<0.001, Fig S1D^i-iv^, S1E). These results suggest that Gsx2 is the main regulator of *dlx6a* in the telencephalon, and in the diencephalon neither Gsx1 nor Gsx2 are essential for *dlx6a* expression.

We also assessed whether Gsx1 or Gsx2 regulate expression of *forkhead box P2* (*foxp2*) in zebrafish. *foxp2* is a gene belonging to the Forkhead domain transcription factors, which are an evolutionarily conserved group of proteins that have roles in early developmental patterning^74^. In humans, *FOXP2* is critical for speech and language development^58^; mutations in *FOXP2* lead to poor linguistic and grammatical skill development and abnormal control of facial movements^75^. *foxp2* is expressed in the nervous system in zebrafish in many overlapping brain regions which we report *gsx1* and *gsx2* expression in, including the telencephalon, diencephalon, optic tectum, hindbrain, and spinal cord^60^. Thus, we were interested in determining if *foxp2* is regulated by either Gsx1 or Gsx2, particularly in the forebrain. Zebrafish *foxp2* possesses putative Gsx1 and Gsx2 enhancers, as well as a putative enhancer sequence that both Gsx1 and Gsx2 could bind to (Fig S1A). Although *foxp2* expression was not statistically different amongst genotypes (p=0.312, Fig S1F^i-iv^, S1G), it is reduced to some degree in *gsx1Δ11*+/+;*gsx2Δ13*−/− and *gsx1Δ11*−/−;*gsx2Δ13a*−/− embryos. This indicates that Gsx2 partially regulates *foxp2* expression in the telencephalon specifically at this age.

## DISCUSSION

### *gsx1* and *gsx2* expression during neurodevelopment in zebrafish embryos and larvae

In this study, we comprehensively document *gsx1* and *gsx2* expression in embryonic and larval zebrafish using multiple strategies, and our analysis presents a time-course for their co-expression during neurodevelopment. In embryonic and larval stages in zebrafish, *gsx1* is expressed in the diencephalon, hypothalamus, preoptic region, hindbrain, cerebellar plate, spinal cord, optic tectum, and pretectum. Across these ages *gsx2* is expressed in the telencephalon, hypothalamus, pallium, olfactory bulb, and hindbrain. These patterns are largely consistent with expression of *Gsx1* and *Gsx2* in mouse^24,29^, medaka^26^, *Xenopus*^25^, and previous reports in zebrafish^10,27,30^ with minor exceptions. In *Xenopus*, *Gsx2* is first detected slightly earlier than *Gsx1*, however we report in zebrafish that *gsx1* is expressed at 10 hpf slightly earlier than *gsx2* at 12 hpf. Furthermore, we report that *gsx2* and not *gsx1* is expressed in the olfactory bulb in zebrafish, however in *Xenopus Gsx1* and not *Gsx2* is expressed in this region.

Prior to our study, a comprehensive knowledge of the unique and overlapping roles for Gsx1 and Gsx2 across the vertebrate brain was not attainable due to the lack of gene expression data. FISH revealed that regional co-expression of *gsx1* and *gsx2* occurs in the hindbrain, hypothalamus, and preoptic area in zebrafish, however they rarely co-localize in the same cells. In the hindbrain *gsx1* and *gsx2* exist in two adjacent dorsolateral columns, with *gsx2* dorsal to *gsx1*, consistent with previous reports and their roles in dorsoventral patterning^24,25^. In mouse, *Gsx1* regulates cerebellar neuronal progenitor identity through a temporally-regulated BMP/SMAD signaling gradient^46,47^, and in zebrafish *gsx2* is reported to specify neuronal fate in the inferior olivary nuclei of the medulla^30^. Outside of these studies the coordinate roles for Gsx1 and Gsx2 in the hindbrain remain under studied. Our findings demonstrate that expression of *gsx1* and *gsx2* remain distinct from each other dorsoventrally, and starting at 48 hpf expression of *gsx1* begins to extend ventrally while *gsx2* is restricted dorsally. These patterns may represent the outgrowth of axons from maturing neuronal progenitors, which would agree with previously reported roles for *Gsx2* and *Gsx1* in regulating progenitor proliferation and differentiation, respectively^22^. Thus, this work provides an essential foundation for future studies to interrogate the functional roles of Gsx1 and Gsx2 in the hindbrain.

*gsx1* expression in the hypothalamus has been shown in medaka, *Xenopus*, mice, and zebrafish^24–27,41^, however no roles for *gsx2* in the hypothalamus have been reported. We show that *gsx1* and *gsx2* are regionally co-expressed in the hypothalamus in zebrafish, which necessitates further studies of Gsx2 function in this region. Zebrafish *gsx2* expression in the hypothalamus begins between 24-30 hpf (Fig 2J, 2L), slightly earlier than the onset of *gsx1* in this region between 30-48 hpf (Fig 2E, 2G). Expression of both *gsx1* and *gsx2* is sustained in the hypothalamus through 6 dpf (Fig 6). Functions for Gsx1 and Gsx2 in this region in zebrafish could be similar to reports in mouse forebrain showing that *Gsx2* maintains neuronal progenitor pools and *Gsx1* drives neuronal differentiation^22^. Interestingly, one single-cell sequencing report conducted in the adult (1-2 year) zebrafish hypothalamus reported Gsx1 as a transcription factor significantly associated with 13 genes categorized as either neuropeptide, neurotransmitter, ion channel, or synaptic genes^76^. Identification of Gsx1 as an important regulatory factor in the mature zebrafish hypothalamus suggests prolonged requirements for Gsx1 in hypothalamic function. As our *gsx1* mutant zebrafish survive through adulthood, roles for Gsx1 in the development and function of the hypothalamus along with associated growth, behavioral, and metabolic changes can be investigated in the future.

### Mutations in *gsx1* and *gsx2* in zebrafish disturbs early growth and development

We observed a reduced growth phenotype in *gsx1* mutant zebrafish through adulthood. These studies provide a detailed description of the onset of significant growth deficits as well as the basic trend and continuation of these deficits. We observed that significant deficits were not present at 4 dpf, but appeared by 14 dpf, allowing us to determine the relative window under which these deficits begin. This data is consistent with work in *Gsx1* mutant mice, which were the same size as their wild-type siblings at birth but began to show growth deficits as development progressed^41^. Unlike *Gsx1* mutant mice, our *gsx1* mutant zebrafish survive to adulthood, permitting investigations of later Gsx1 function. The premature death of *Gsx1* mutant mice is largely attributed to defects in forebrain neurogenesis and disruptions in ascending cortical interneuron migration^22,44,51^, thus continued examination of the impact of mutations in *gsx1* in zebrafish will further elucidate its important neurodevelopmental and later roles in vertebrates.

We have additionally identified a unique swim bladder inflation failure phenotype in *gsx2* mutant zebrafish that prevents their survival under standard rearing conditions, supporting the critical role for Gsx2 in growth and development amongst vertebrates. *Gsx2* mutant mice fail to survive more than a day following birth, however also exhibit severely disrupted forebrain and hindbrain morphology. Comprehensive knowledge of GSX1 and GSX2 function together and separately is minimal outside of the mouse forebrain^34,44,51^ and few reports in the cerebellum^30,46,77^ and spinal cord^7,10,48^. As such, analysis of Gsx1 and Gsx2 function in our zebrafish mutants in these and other CNS regions can supplement these reports.

### Identifying Gsx1 and Gsx2 target genes in the zebrafish forebrain

We demonstrated differential regulation of *dlx2a*, *dlx2b*, *dlx5a*, *dlx6a*, and *foxp2* by Gsx1 and Gsx2 in the forebrain of our *gsx1* and *gsx2* mutant zebrafish. A complex relationship between the *Gsx* and *Dlx* genes has been reported in the mouse forebrain^51^ that facilitates regulation of a major transcriptional control program dictating the expression of diverse target genes. The *Dlx* pathways in the forebrain also serve to regulate the differentiation of inhibitory projection neurons and interneurons that migrate to mature regions like the cortex and olfactory bulb^53–55^. Conservation of the Gsx/Dlx regulatory network in zebrafish is significant in that it establishes initial understanding of Gsx function in neurodevelopment in zebrafish. Our embryonic and larval stage whole brain expression analyses also justify continued investigations of Gsx function together and separately across brain regions to add to our knowledge of their role in neurodevelopment across vertebrates.

Our data suggests that in zebrafish, Gsx2 is largely responsible for regulating expression of *dlx2a* and *dlx2b*. Through WISH we identified significant reductions in *dlx2a* and *dlx2b* expression in the telencephalon of *gsx1Δ11*+/+;*gsx2Δ13a*−/− and *gsx1Δ11*−/−;*gsx2Δ13a*−/− embryos (Fig 8). In the diencephalon, *dlx2a* expression was reduced in both *gsx1Δ11*+/+;*gsx2Δ13a*−/− and *gsx1Δ11*−/−;*gsx2Δ13a*−/− embryos, however *dlx2b* expression was only reduced in *gsx1Δ11*−/−;*gsx2Δ13a*−/− embryos, suggesting that Gsx1 may be compensating for Gsx2 and sustaining *dlx2b* expression in the diencephalon specifically. RT-qPCR analysis of *dlx2b* expression was consistent with these results, revealing significant reductions in *gsx1Δ11*+/+;*gsx2Δ13a*−/− embryos and more significant reductions in *gsx1Δ11−/−;gsx2Δ13a−/−* embryos. Unlike our WISH analysis, RT-qPCR shows that *dlx2a* expression is only significantly reduced in *gsx1Δ11*−/−;*gsx2Δ13a*−/− embryos and not *gsx1Δ11*+/+;*gsx2Δ13*−/− embryos. One potential explanation for this variability is the pattern in which *dlx2a* expression is reduced by WISH. In *gsx1Δ11*−/−;*gsx2Δ13a*−/− embryos, *dlx2a* expression is lost in the telencephalon, where *gsx2* is expressed, however not in the diencephalon, where *gsx1* is expressed. This variability could also be related to alternative transcript detection through qPCR. However, our *in situ* probe for *dlx2a* detects a product that overlaps completely with the transcript amplified by our *dlx2a* qPCR primers, and we predict these targets are identical for both *dlx2a* splice variants that exist in zebrafish.

Expression of zebrafish *dlx5a* is largely regulated by Gsx2 in the telencephalon, as we identified significant reductions in *gsx1Δ11*+/+;*gsx2Δ13a*−/− and *gsx1Δ11*−/−;*gsx1Δ11*−/− embryos (Fig S1B, S1C). However, in the diencephalon it appears that both Gsx1 and Gsx2 regulate *dlx5a* expression, as significant reductions were only identified in *gsx1Δ11*−/−;*gsx2Δ13*−/− embryos. In turn, this suggests that Gsx1 is in part compensating for loss of Gsx2 function in *gsx1Δ11*+/+;*gsx2Δ13a*−/− embryos, which were not significantly reduced compared to wild-types. Interestingly, expression of *dlx6a* appears to be regulated most strongly by Gsx2 in the telencephalon only, agreeing with initial predictions based on enhancer sequence presence (Fig S1A, S1D, S1E). Collectively, these results demonstrate that a complex relationship between the *gsx* and *dlx* genes exists in zebrafish that is reminiscent of reports in other vertebrates^51^. Future studies will focus on confirming more Gsx1 and Gsx2 target genes in zebrafish in order to elucidate their unique and overlapping roles during CNS development.

Outside of confirming that several *dlx* orthologs are regulated by Gsx1 and Gsx2 in the zebrafish forebrain, we also found that although not statistically significant, *foxp2* is weakly regulated in the telencephalon by Gsx2. It is important to note that this study was conducted at 30 hpf only, and *foxp2* expression begins in zebrafish as early as 10 hpf in the presumptive forebrain^60^ and is documented through 3 months of age^78^. The onset of *foxp2* expression is similar to the onset of *gsx2* in the presumptive forebrain at 12 hpf (Fig 2H), however during neurodevelopment *foxp2* is expressed in several overlapping regions with the *gsx* genes, such as the optic tectum, hindbrain, and spinal cord. Minimal regulation by Gsx2 in the telencephalon is consistent with a requirement for Gsx2 in early forebrain patterning observed throughout these experiments, particularly in the telencephalon. It is interesting to note that expression of *foxp2* and *dlx6a* minimally overlaps in dorsal subpallial regions in zebrafish^60^, and *dlx6a* is regulated by Gsx2 in this dorsal telencephalic region only. Collectively, our approaches for identifying and validating target genes for Gsx1 and Gsx2 during neurodevelopment provide a new *in vivo* model for gaining even greater insight into regulatory roles of these and other transcription factors across CNS gene networks.

## ACKNOWLEDGEMENTS

The authors would like to thank Timothy Driscoll, PhD and Jessica Towey (WVU Dept of Biology) for advice and equipment training and use related to the RT-qPCR experiments. Benjamin Feldman, PhD, director of the NICHD Zebrafish Core (NIH), provided guidance early on related to TALEN mutant generation before CRISPR/Cas9 became all the rage. Eric Horstick, PhD and Andrew Dacks, PhD (WVU Dept of Biology) carefully read the manuscript and provided valuable preliminary feedback.

## CONTRIBUTIONS TO THE MANUSCRIPT

SAB and RAC designed and performed experiments, wrote and edited the manuscript, analyzed and interpreted the results; EIS, ZAD, SNP, ARA, LCF, and RLP performed and assisted with experiments, collected data, and ran initial analyses.

---

**Figure S1.**
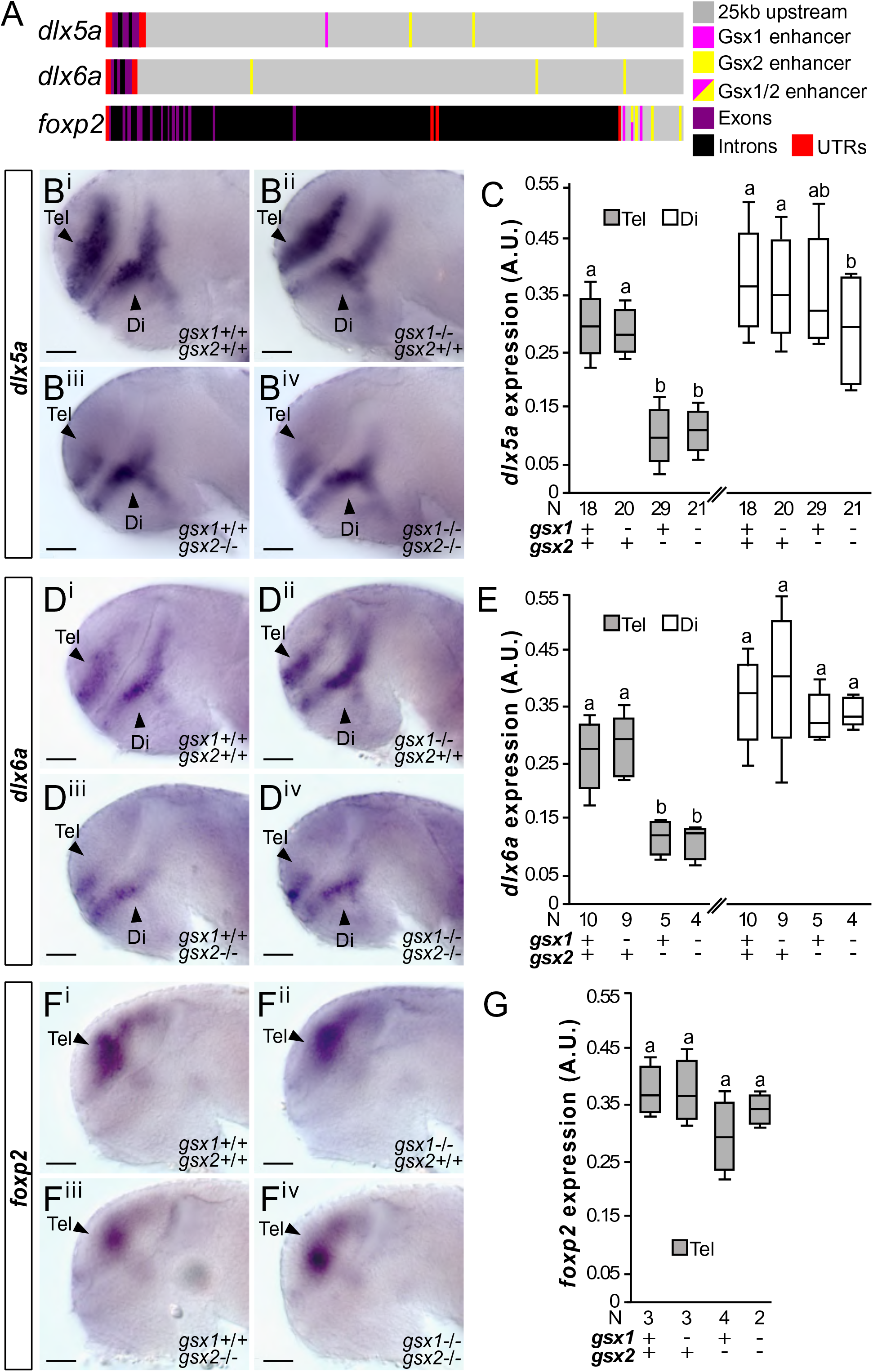
Regulation of *dlx5a, dlx6a*, and *foxp2* by Gsx1 and Gsx2. **A)** Schematic of Gsx1 and 2 enhancer sequences upstream of the zebrafish *dlx5a*, *dlx6a*, and *foxp2* gene bodies. **B^i^-B^iv^.** *dlx5a* expression at 30 hpf in wild-type, *gsx1Δ11*−/−, *gsx2Δ13a*−/−, and *gsx1Δ11*−/−;*gsx2Δ13a*−/− zebrafish. Images are 20X compound scope images with samples mounted under cover glass, eyes dissected, and anterior facing left. Scale bar = 50μm. **C.** FIJI-ImageJ quantification of *dlx5a* expression. Different letters represent significant differences. **D^i^-D^iv^.** *dlx6a* expression at 30 hpf in wild-type, *gsx1Δ11*−/−, *gsx2Δ13a*−/−, and *gsx1Δ11*−/−/*gsx2Δ13a*−/− zebrafish. **E.** Quantification of *dlx6a* expression. **E^i^-E^iv^.** *foxp2* expression at 30 hpf in wild-type, *gsx1Δ11*−/−, *gsx2Δ13a*−/−, and *gsx1Δ11*−/−/*gsx2Δ13a*−/− zebrafish. **G.** Quantification of *foxp2* expression area.

**Table S1.** List of NCBI sequences for gene and protein alignments and analyses.

| Gene sequences |  |  |
| --- | --- | --- |
| Species | Gene | Accession number |
| *Danio rerio* | *gsx1* | NM_001012251.1 |
| *Danio rerio* | *gsx2* | NM_001025512.2 |
| *Danio rerio* | *dlx2a* | AF349437.2 |
| *Danio rerio* | *dlx2b* | NM_131297.2 |
| *Mus musculus* | *Dlx2* | NM_010054.2 |
| *Homo sapiens* | *Dlx2* | NM_004405.4 |
| *Danio rerio* | *dlx5a* | NM_131306.2 |
| *Danio rerio* | *dlx6a* | NM_131323.1 |
| *Danio rerio* | *foxp2* | NM_001030082.2 |
| Protein sequences |  |  |
| Species | Protein | Accession number |
| *Danio rerio* | Gsx1 | AAI65050.1 |
| *Mus musculus* | Gsx1 | AAI37770.1 |
| *Rattus norvegicus* | Gsx1 | NP_001178592.1 |
| *Homo sapiens* | Gsx1 | NP_663632.1 |
| *Xenopus tropicalis* | Gsx1 | NP_001039254 |
| *Oryzias latipes* | Gsx1 | NP_001098303 |
| *Lepisosteus oculatus* | Gsx1 | XP_006627824 |
| *Danio rerio* | Gsx2 | AAI64330.1 |
| *Mus musculus* | Gsx2 | NP_573555.1 |
| *Rattus norvegicus* | Gsx2 | NP_001131035.1 |
| *Homo sapiens* | Gsx2 | NP_573574.2 |
| *Xenopus tropicalis* | Gsx2 | AAI58504.1 |
| *Oryzias latipes* | Gsx2 | NP_001116381 |
| *Lepisosteus oculatus* | Gsx2 | XP_006630061 |
| *Drosophila melanogaster* | Ind | NP_996087.2 |

**Table S2.**
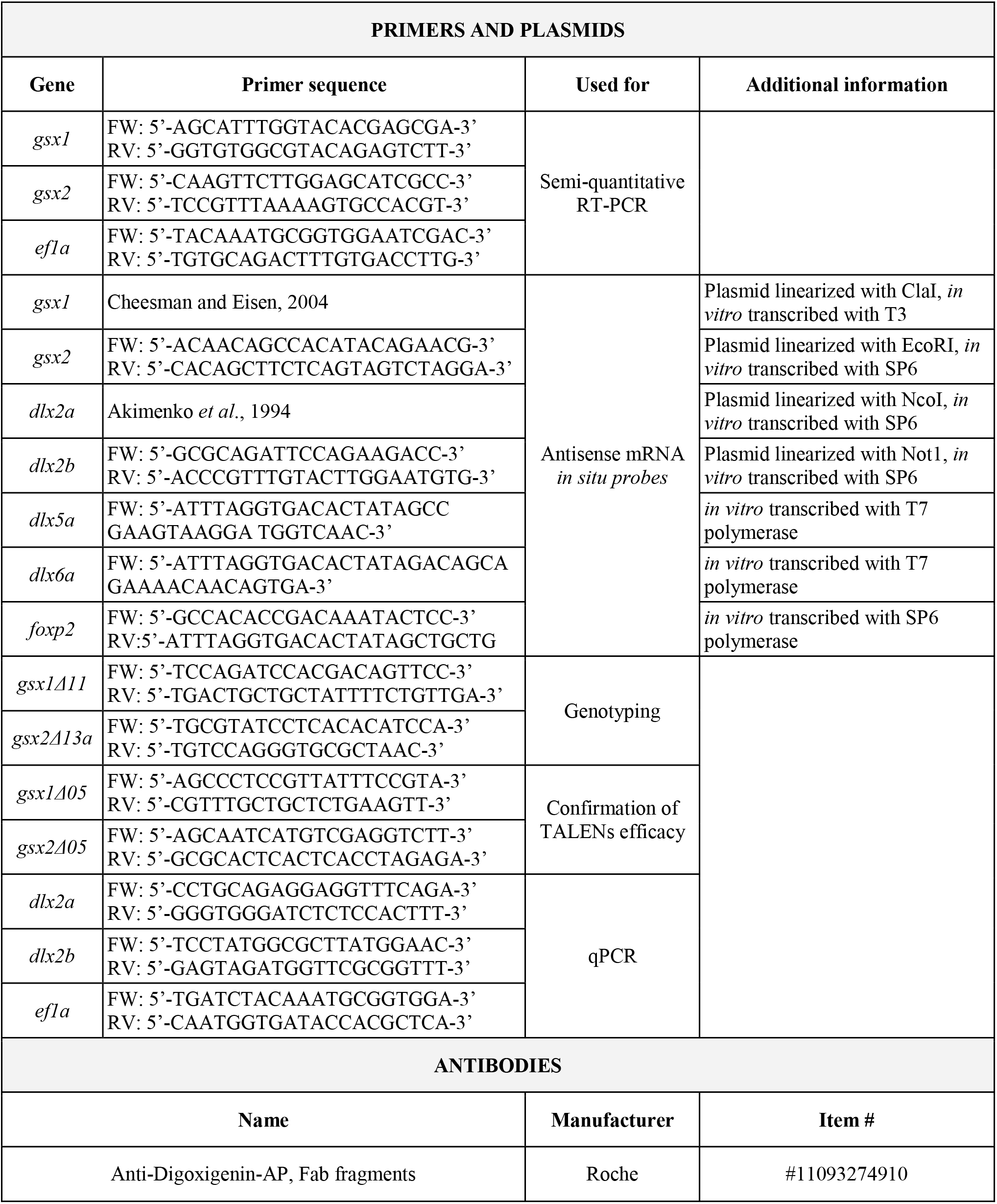
Table of primers, plasmids, and antibodies.

